# Extremely low effective population size in a captive-bred population: partial mitigation through management practices

**DOI:** 10.64898/2026.05.12.724519

**Authors:** Amaïa Lamarins, Robin S. Waples, Jorma Piironen, Craig R. Primmer

**Affiliations:** Department of Organismal and Evolutionary Biology, Faculty of Biological and Environmental Sciences, University of Helsinki, Helsinki, Finland; School of Aquatic and Fishery Sciences, University of Washington, Seattle, Washington, USA; Natural Resources Institute Finland (Luke), Natural Resources, Migratory Fish and Regulated Rivers, Joensuu, Finland

**Keywords:** Atlantic salmon, captive-bred, effective population size, landlocked, management, Saimaa

## Abstract

Effective population size (*N*_*e*_) is a critical parameter for evaluating the evolutionary and persistence potential of endangered populations and for designing sustainable conservation strategies. Captive breeding and release programs are widely used across taxa to reduce risk of extinction when natural reproduction is insufficient or no longer possible, making it essential to assess their consequences. We used the case study of the landlocked Saimaa salmon (*Salmo salar*), one of the most critically en-dangered salmonid populations in Europe, with unique evolutionary significance due to its isolation from other populations since the last glaciation. Using long-term demographic data (1969-2024) from wild-caught founders of a captive breeding and release program, we estimated the effective population size under multiple scenarios of variance in reproductive success. Across scenarios, *N*_*e*_ ranged from 33 to 81 individuals, representing 32%-75% of the census size. Captive breeding practices aimed at equalizing parental contributions during fertilization and early life stages increased *N*_*e*_ by 12% compared to natural reproductive conditions. However, variation in survival after early developmental stages, typically beyond direct management control, remained a key determinant of *N*_*e*_. Despite recent increases in the number of founders, the population remains genetically vulnerable due to historical bottlenecks. These results highlight that while captive breeding programs can partially mitigate genetic risks, their effectiveness depends critically on both controlled and uncontrolled sources of variance in reproductive success. Strengthening such programs may require combining breeding management with habitat restoration and, where appropriate, genetic rescue to ensure the long-term evolutionary potential of such unique and endangered populations.

## 2 Introduction

Monitoring the effective population size (*N*_*e*_) of endangered populations is critical for evaluating their evolutionary and persistence potential and for designing sustainable breeding and conservation strategies. *N*_*e*_ determines the rate of genetic diversity loss and inbreeding increase in a population, and it constrains selection and adaptation through genetic drift. Effective population size is primarily influenced by the number of breeding individuals, but also by their relative reproductive success, and is thus sensitive to variation in reproductive success and sex ratio (Wright, 1938; Frankham, 1995; Waples, 2025). Demographic information can provide valuable insights into the realized effective population size in small, intensively managed populations where genetic sampling is limited, but demographic details and knowledge of management practices are plentiful. Previous studies have shown that estimates derived from genetic and demographic data are often similar (Saura et al., 2008; Hoehn et al., 2012; Serbezov et al., 2012; Perrier et al., 2014).

Captive breeding and release programs are widely used across taxa to reduce risk of extinction when natural reproduction is insufficient or no longer possible. These interventions have been implemented in a broad range of endangered species, including fishes, amphibians, birds, and mammals (Fraser, 2008; Harding et al., 2016; Redford et al., 2011). In some cases, populations have become entirely dependent on continuous captive propagation for their persistence in the wild. Well-known examples include the California condor (*Gymnogyps californianus*), which was reduced to a fully captive population before reintroduction (Bakker et al., 2024), the black-footed ferret (*Mustela nigripes*), which persists through intensive captive breeding and release efforts (Santymire et al., 2014), and several salmonid populations maintained through long-term hatchery supplementation (Fraser, 2008). While such programs can reduce the risk of immediate extinction, they often create dependence on human intervention and may alter demographic structure, reproductive variance, and genetic composition (Snyder et al., 1996; Araki et al., 2007*a*). Assessing the demographic and genetic consequences of long-term captive breeding and release programs and their practices is therefore critical for evaluating the sustainability and evolutionary potential of managed populations.

Here, we use the landlocked Atlantic salmon (*Salmo salar* m. *sebago*) inhabiting the Lake Saimaa system in eastern Finland as a case study for such a scenario. Saimaa salmon represent one of the most critically endangered salmonid populations in Europe (Hyvärinen et al., 2019). Isolated from anadromous populations of the Baltic Sea basin after the last glaciation approximately 10,000 years ago, the Saimaa salmon evolved as a distinct non-anadromous landlocked lineage within the Vuoksi watercourse (Lumme et al., 2015). However, extensive human modifications of river systems during the 20th century, including dam construction and hydropower development, have destroyed almost all natural breeding habitats and restrict access to remaining breeding and nursery habitats. In particular, the construction of the Kuurna dam in 1971 blocked salmon migration routes from the lake Saimaa to the main breeding rivers, leading to the collapse of natural reproduction (Pursiainen et al., 1998).

Because of these anthropogenic changes, the persistence of Saimaa salmon in the wild has relied almost entirely on captive breeding and artificial propagation since the 1960s (Kallio, 1986; Pursiainen et al., 1998; Kaijomaa et al., 2003). Since 1969, mature individuals ascending below hydropower dams have been captured annually for founding captive-reared breeding stocks used for juvenile releases into the lake system. However, survival during the lake phase remains extremely low due to post-release mortality, incidental by-catch in fishing nets, and targeted trolling (Pursiainen et al., 1998; Kaijomaa et al., 2003; Huuskonen et al., 2007). As a result, the number of wild-caught individuals founding the breeding stocks has been very low, averaging around 10 individuals per year during the early decades of the program (1980s). These demographic constraints, combined with the closed captive-bred system, have led to severe genetic bottlenecks and reduced genetic diversity (Koljonen et al., 2002; Säisä et al., 2005; Tonteri et al., 2005; Koljonen et al., 2022). Evidence of inbreeding depression has been reported, such as developmental deformities in captive-reared juveniles (Tiira et al., 2006). Despite continuous breeding in captivity and complete fishing ban on wild Lake Saimaa salmon since 2016, the long-term viability of this unique population remains uncertain. Therefore, there is a need for improved management practices and a deeper understanding of the population’s genetic and demographic constraints.

In this study, we used historical data on founders composition, age structure, and fecundity dating back to the 1970s to estimate the effective population size of the Saimaa salmon under its current management regime and alternative assumptions. Our approach used demographic modeling of *N*_*e*_ by considering likely variation in reproductive success among individuals. Because some information was incomplete, we combined available data with expert knowledge to explore a range of plausible scenarios of variance in reproductive success. Specifically, we evaluated four contrasting scenarios reflecting increasing levels of variance in fecundity, difference in survival processes (random vs family-correlated), and the effect of captive breeding practices that reduce variance in reproductive success. We expect that increasing variance in reproductive success via fecundity or family-correlated survival will lead to substantial reductions in *N*_*e*_, whereas management practices may mitigate these effects. Beyond this specific case study, such an approach can be broadly applied to other endangered populations managed through captive breeding and release programs, particularly where detailed demographic data are available but genetic information is limited. This framework provides a valuable tool for understanding the genetic consequences of management practices and for guiding future conservation actions aimed at maintaining the long-term evolutionary potential of such unique and isolated populations.

## 3 Methods

### 3.1 Study system and available data

We use the landlocked Atlantic salmon population of the Lake Saimaa system in eastern Finland as a case study (Fig. 1). Since the collapse of natural reproduction, mature individuals ascending rivers have been captured to found captive breeding stocks and for subsequent juvenile release. The first breeding stocks were founded at the Kontiolahti hatchery, located 2 km from the Kuurna hydropower dam. Cultivation of landlocked salmon was then continued at the Enonkoski Aquaculture Station of the Natural Resources Institute Finland since 1983. Wild adults ascending below the Kuurna hydropower dam on the River Pielisjoki have been captured annually since 1969 to found captive breeding stocks, with additional founders collected from the River Lieksanjoki since 1992 and from the Kermankoski rapids since 1995. Fertilized eggs are reared under controlled captive conditions, and juveniles are mainly released as two-years old smolts into multiple lake areas within the watershed. After release, individuals undergo a natural growth phase in the lake before returning as adults, at which point a subset is again collected for captive breeding. The population thus forms a closed, captive-bred system with limited natural reproduction. All procedures were performed in compliance with relevant laws and institutional guidelines.

**Figure 1:**
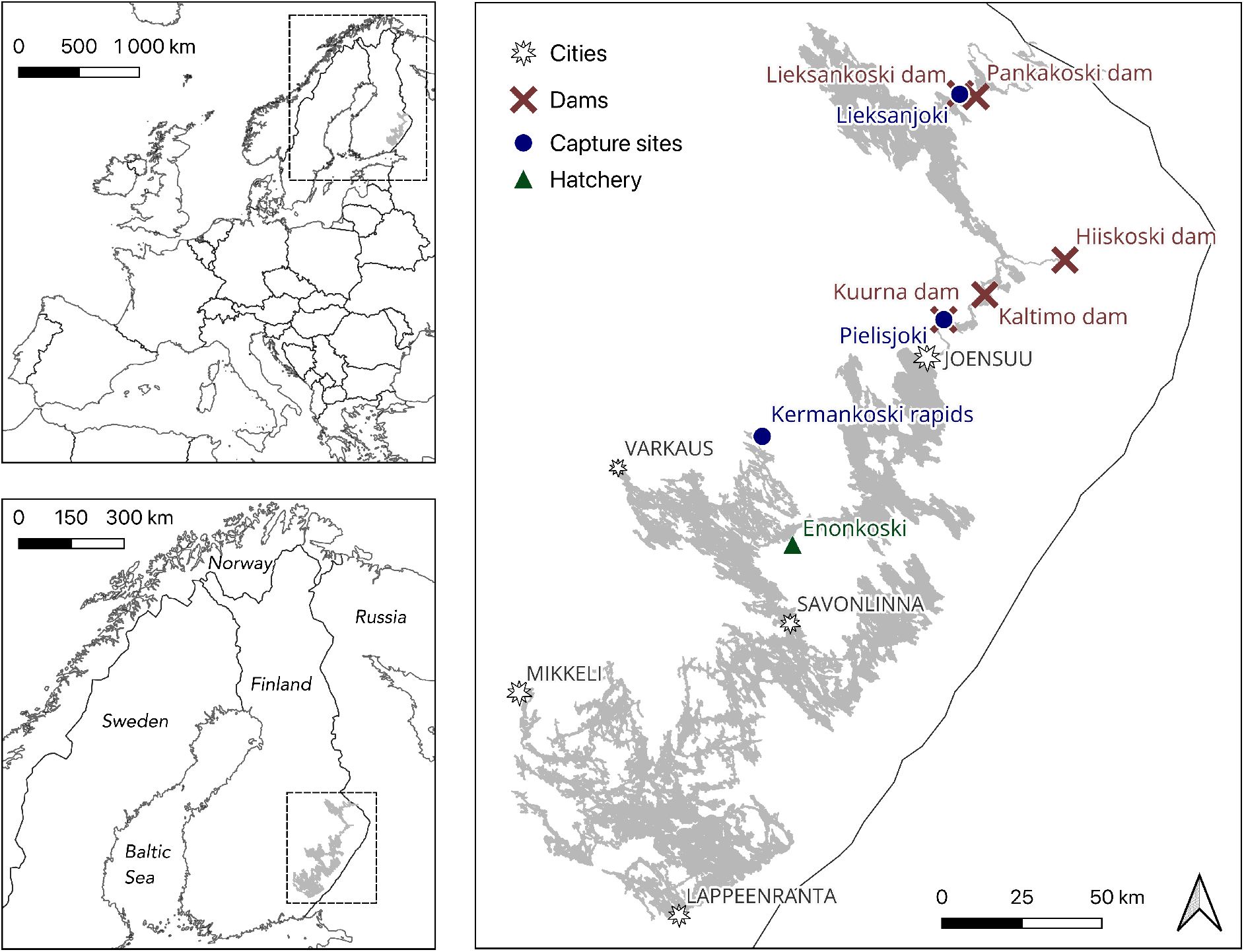
Map of the Lake Saimaa salmon system (grey area) in eastern Finland, showing the locations of dams (crosses), breeders capture sites (circles), and the hatchery (triangle). Source data: Finnish Environment Institute (Syke).

The data used in this study consist of annual records of the number of wild-caught individuals used for founding breeding stocks between 1969 and 2024 from the Pielisjoki River, the Lieksanjoki River and the Kermankoski rapids. For the period 1969-1974, the number of founders was estimated. The number of females was inferred from stripped egg volumes using the linear relationship between egg volume and number of females observed during 1975-1985, assuming a similar life-history composition. The number of males during 1969-1974 was estimated based on the observed sex ratio during 1975-1985. From 1975 to 2024, direct counts of breeding stocks founders were used in the analyses.

Monitoring programs also provided information on the age distribution by sex of wild-caught breeders, allowing estimation of the expected proportion of females and males maturing at each age between 4+ and 7+ years (corresponding to 2–5 years spent in the lake). Male typically mature at ages 5-6 after 3-4 years in the lake, whereas females generally mature at ages 4-5 after 2-3 years in the lake (Pursiainen et al., 1998; Kaijomaa et al., 2003). We then estimated the generation length, from the birth of parents to the birth of their offspring, at 6.43 years (Table S1).

Average female fecundity (i.e., number of eggs per female) was estimated for each age group based on mean body weight and an empirical relationship between female weight and egg number. These estimates, provided by hatchery staff, were approximately 3500, 4300, 5200 and 7300 eggs for females of age 4+, 5+, 6+ and 7+, respectively. Sperm availability was assumed not to limit fertilization, as matrix fertilization designs have been used since the early 1980s, ensuring that each male fertilizes eggs from multiple females. Therefore, mean reproductive success among males was assumed to be approximately equal across individuals and age classes. In a small number of years, additional gametes of some males and females from previously created breeding stocks were used in later years but were not considered in *N*_*e*_ estimates for simplicity.

### 3.2 Demographic estimation of effective population size

We estimated the effective population size (*N*_*e*_) following the framework of Waples (1990, 2002*a*, 2025). This approach quantifies how unequal reproductive contributions among individuals reduce the genetic diversity relative to census population size, using demographic information on sex ratio, age structure, mean and variance in reproductive success. Annual sex-specific effective numbers of breeders (inbreeding *N*_*bI,s*_ or *N*_*b,s*_) were first calculated based on mean and variance in reproductive success, and then combined across sexes to obtain an overall annual *N*_*b*_. Assuming semelparity, the generational effective size *N*_*e*_ is then derived from the harmonic mean of *N*_*b*_ across years and the generation length.

#### 3.2.1 Generational effective population size

Following Waples (1990, 2002a, 2006), and assuming semelparity, the effective population size per generation *N*_*e*_ was derived by multiplying the generation time *g* by the harmonic mean of the overall effective number of breeders *N*_*b*_(*t*) across years:

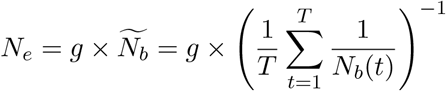

where *T* is the number of years considered, and *g* the generation length. The generation length corresponds to the average age of parents producing eggs, weighted by age structure, and was estimated here at g=6.43.

The overall effective number of breeders, *N*_*b*_(*t*), is adjusted for sex ratio and computed for each year following Wright (1938):

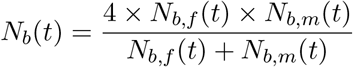

where *N*_*b,f*_ (*t*) and *N*_*b,m*_(*t*) are the effective numbers of breeding females and males, respectively, for the year *t*.

The effective number of breeders and thereby the effective population size can be calculated at different time of sampling and thus life stages, such as the egg stage or the adult stage. *N*_*e*_ can also be estimated over a full life cycle according to two alternative assumptions: i) random survival, where all families experience equal survival probability from the sampling time (e.g., eggs), ii) non-random survival, where survival is family-dependent (some families survive entirely and others are lost).

#### 3.2.2 Effective number of breeders for i) random survival

For each year *t*, the actual number of wild-caught breeding females *N*_*f*_ (*t*) and males *N*_*m*_(*t*) were distributed among four age classes (*a* = 1,.. ., 4) according to the observed proportions of maturing adults (*p*_*s,a*_, where *s ∈ {f, m}* for females and males, respectively). The expected number of breeders per age class was thus:

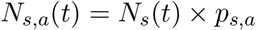

The sex-specific effective number of breeders (inbreeding *N*_*bI,s*_ or *N*_*b,s*_) was computed following Lande and Barrowclough (1987):

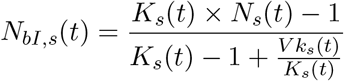

where *K*_*s*_(*t*) is the mean number of offspring per individual and *V k*_*s*_(*t*) is its associated variance.

These were obtained as:

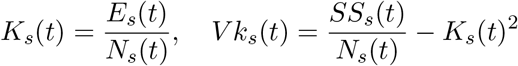

where *E*_*s*_(*t*) and *SS*_*s*_(*t*) are the sum across all ages of *E*_*s,a*_, the expected total number of eggs produced by individuals of age *a*, and *SS*_*s,a*_, the sum of squares of individual reproductive success by age *a*:

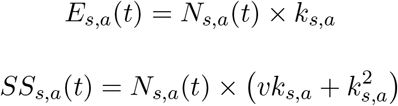

Here, *k*_*s,a*_ and *vk*_*f,a*_ denote the mean and variance in fecundity for age class *a*.

For males, sperm was not considered limiting and the mean fecundity per age class *a* was assumed constant and scaled to the total egg production:

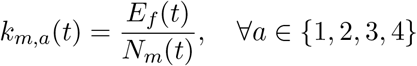

#### 3.2.3 Effective number of breeders for ii) non-random survival

The previous approach assumes random survival from egg to adult, which may be unrealistic if survival differs among families, as pointed out in Moyer et al. (2007). To address this, we applied an alternative method developed by Crow and Morton (1955) and described in Waples (2002*b*). This method is based on the assumption that entire families either survive or not.

To account for the potential non-random survival of families after the egg stage, we considered a fixed proportion of families that successfully survived after the egg stage, denoted *ω*. Under this assumption, the number of surviving individuals of age *a* and sex *s* was:

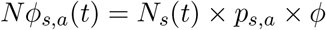

Assuming a constant population size, the mean reproductive success of surviving individuals was:

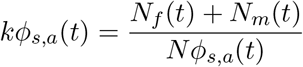

The variance in reproductive success is then modified to account for non-random survival following Crow and Morton (1955) approximation for random family survival:

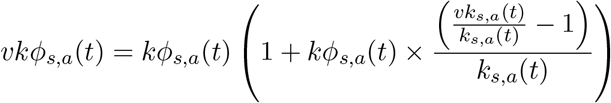

where *k*_*s,a*_(*t*) and *vk*_*s,a*_(*t*) are the original mean and variance in reproductive success before family-dependent survival.

Subsequent steps for computing *SS*_*s*_(*t*), *K*_*s*_(*t*), *V k*_*s*_(*t*) and *N*_*b,s*_(*t*) followed the same approach as for the random survival case. These two approaches represent alternative extremes - complete random survival vs complete family-correlated survival - and thus provide a plausible range for the effective population sizes under different degrees of family-level selection.

### 3.3 Scenarios of variance in reproductive success

Because direct observations of variance in reproductive success were unavailable, we evaluated a set of alternative scenarios reflecting natural variation and captive breeding and rearing practices.

#### 3.3.1 S1: Natural variation in sex ratio and fecundity

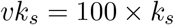

This baseline scenario assumes moderate natural variation in fecundity (Moffett et al., 2006; de Eyto et al., 2015; Maamela et al., 2025) within age classes for both sexes, representing conditions without intentional family-size equalization. It also assumes random survival from egg to adult.

#### 3.3.2 S2: Higher variance in reproductive success

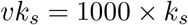

This scenario represents stronger variance in reproductive success (Heinimaa and Heinimaa, 2004), which may arise from unequal fertilization success or egg viability, potentially increased in captive-reared breeding stocks. It provides an upper bound on reproductive variance expected at the egg stage under natural conditions with limited control over fertilization or early survival.

However, it still assumes random survival until the adult stage. Consequently, the two above scenarios generally provide upper limits to *N*_*b*_ and *N*_*e*_.

#### 3.3.3 S3: Non-random survival post-egg

Building on the higher-variance assumption (S2), this scenario further considers that survival from the egg to adult stage may be family-correlated rather than random. Such correlations can arise from family-specific egg quality, early growth, or susceptibility to environmental conditions. We modeled this effect by assuming that only a proportion of families successfully survive beyond the egg stage. We assumed an intermediate value of *ϕ* of 0.5.

#### 3.3.4 S4: Captive-breeding and rearing practices

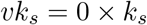

This scenario reflects the breeding and rearing practices applied at the Enonkoski Aquaculture Station of the Natural Resources Institute Finland. The procedure aims to produce breeding stocks with the most even and genetically diverse structure possible. In practice, this is achieved by fertilizing the eggs of each wild-caught female with milt from each male using a so-called fertilization matrix system. Specifically, a small portion of eggs (200-300) from each female is fertilized separately by different males, and the fertilized eggs are incubated in family-specific trays. For example, a 5×5 matrix generates 25 distinct families. After the eggs have developed to eyed stage, an equal number of eggs (usually from 5-10 /family) are sampled from each family to establish a new breeding stock cohort for cultivation. This procedure minimizes family-specific variation and ensures equal genetic contributions among families at this stage. Beyond the eyed-egg stage, family-specific mortality cannot be controlled. The fertilization matrix system, combined with the equalization of family sizes at the eyed-egg stage, effectively minimizes variance in the reproductive success among individuals. Accordingly, we assumed a null variance in fecundity at the eyed-egg stage, followed by non-random survival during subsequent life stages.

## 4 Results

Estimates of the effective number of breeders (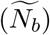), the generational effective population size (*N*_*e*_), and the ratio between the effective and observed number of breeders (*N*_*b*_*/N*) varied among the different scenarios of variance in reproductive success (Figure 2; Table 1). Under the baseline scenario assuming moderate natural variation in fecundity (S1), the estimated effective number of breeders remained below the census size (median *N*_*b*_*/N* = 0.75), with an average effective number of breeders of 12.6, corresponding to an upper limit of *N*_*e*_ of 81 individuals (Figure 2; Table 1). Increasing the variance in reproductive success (S2) slightly reduced both 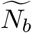 and *N*_*e*_, yielding values of 11.5 effective breeders on average and an effective population size of 74 individuals (9% reduction; Figure 2; Table 1).

**Table 1:** Harmonic mean of annual effective number of breeders 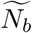, generational effective population size 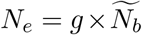, and median of annual ratio between the effective and observed number of breeders *N*_*b*_*/N* during the whole time period, under different scenarios of variance in reproductive success.

| Scenario | $\widetilde{N}_b$ | $N_e$ | $N_b/N$ |
| --- | --- | --- | --- |
| S1 | 12.6 | 81.1 | 0.75 |
| S2 | 11.5 | 73.7 | 0.68 |
| S3 | 5.1 | 33 | 0.32 |
| S4 | 5.7 | 36.9 | 0.36 |

**Figure 2:**
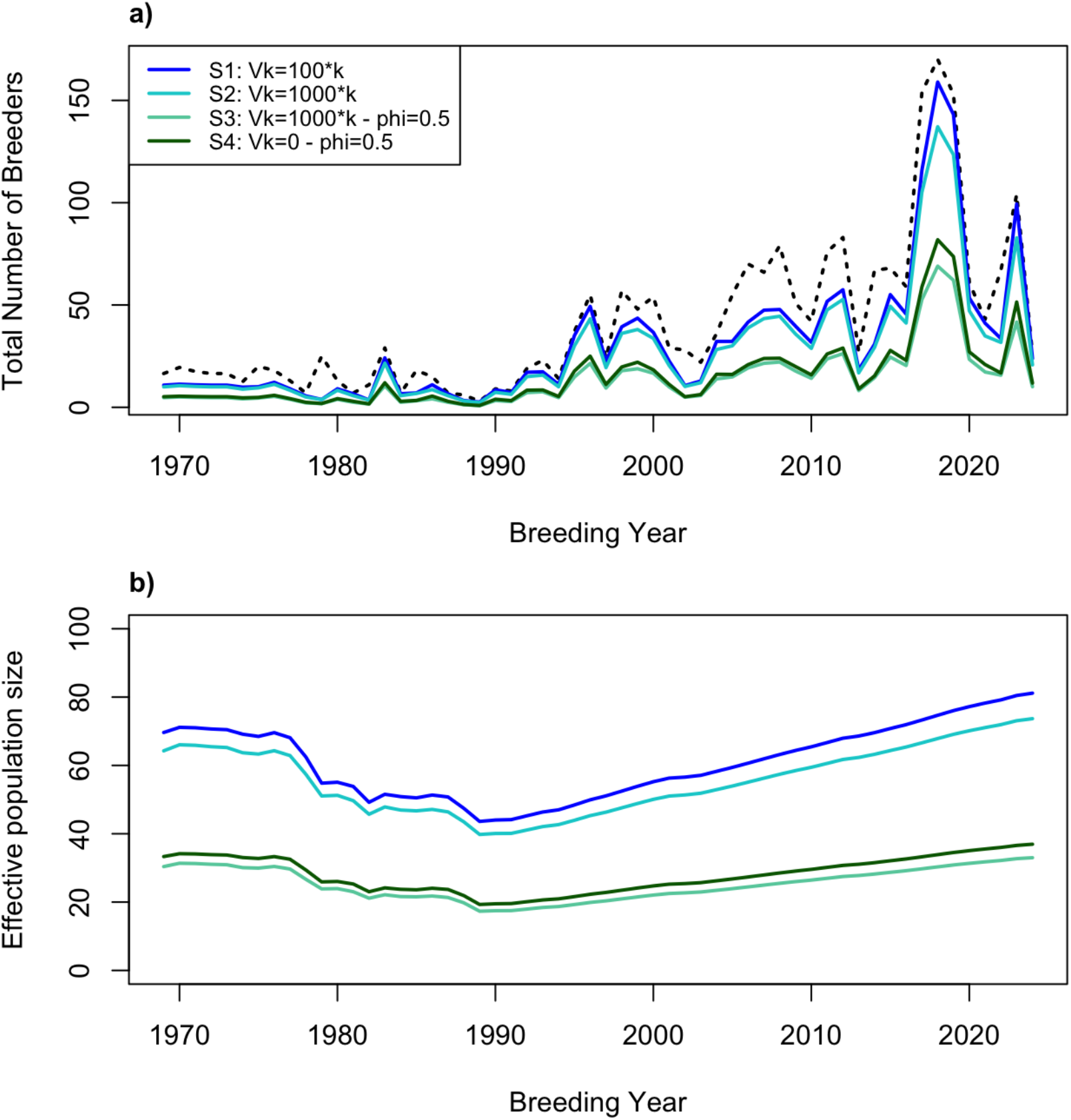
Estimates of the Saimaa salmon annual a) effective number of breeders (*N*_*b*_) and b) generational effective population size (*N*_*e*_) under different scenarios of variance in reproductive success (colored solid lines). Scenarios S1 and S2 assume natural and elevated variance in fecundity, respectively, combined with random survival to the adult stage, and therefore represent upper limits for *N*_*e*_. Scenarios S3 and S4 assume non-random survival until the adult stage, with scenario S4 additionally accounting for captive-breeding practices that reduce variance in fecundity. The total number of wild-caught breeders used in the breeding program is showed by the black dashed line.

Introducing family-correlated survival had a strong effect on reducing the effective population size. When only half of the families survived beyond the egg stage under high reproductive variance (S3), 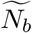 dropped to 5.1 and *N*_*e*_ to 33 individuals (55% further reduction; Figure 2; Table 1). The median ratio between the effective number of breeders and census size also decreased to 0.32 (Table 1). In contrast, the scenario including equalization of family sizes but with the same level of family-correlated survival (S4) resulted in slightly higher values, with a 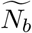 of 5.7, a *N*_*e*_ of 37 individuals (12% increase) and a median *N*_*b*_*/N* ratio of 0.36 (Figure 2; Table 1).

Across all scenarios, the effective population size, after an initial decline prior to the 1990s, gradually increased in parallel with the number of captive breeders (Figure 2). However, despite these increases, all scenarios indicate that the effective size of the Saimaa salmon population has remained critically low.

## 5 Discussion

Captive breeding and release programs are widely used to reduce the risk of extinction when natural reproduction is insufficient. Assessing the demographic and genetic consequences of their practices is critical to ensure the long-term persistence and evolutionary potential of managed populations. Using the landlocked Saimaa salmon population as a case study, we demonstrate how demographic data can be used to estimate effective population size and evaluate the consequences of alternative management scenarios. More broadly, this approach can be applied to other endangered populations where detailed demographic data are available but genetic data may be limited.

As predicted by population genetic theory, the extremely small number of breeders during the early years (1980s–1990s) of the breeding program has led to a severe genetic bottleneck. Our demographic analyses indicate that the effective population size (*N*_*e*_) of the landlocked Saimaa salmon remains critically low under all examined scenarios, despite the long-standing captive breeding program. Across scenarios, *N*_*e*_ ranged from 33 to 81 individuals, which aligns closely with the genetically-based estimate of 48 individuals (95% CI: 32-75) reported by Koljonen et al. (2022). Across our scenarios, the median *N*_*b*_*/N* ratio varied between 0.32 and 0.75. These values are broadly consistent with previous demographic estimates reported for other salmonids populations. For example, Waples (2002a) reported a mean *N*_*e*_*/N* ratio of 0.39-0.61 for a Chinook salmon population, while empirical studies of Steelhead trout found mean *N*_*b*_*/N* ratios of 0.61 (Ardren and Kapuscinski, 2003) and 0.17-0.4 (Araki et al., 2007*b*). Genetic studies of Atlantic salmon populations similarly reported a wide range of *N*_*b*_*/N* or *N*_*e*_*/N* ratios, from 0.12-0.9 (mean 0.36) in Canada (Perrier et al., 2016), 0.18-0.54 and 0.25 in Spain (Consuegra et al., 2005; Saura et al., 2008), and 0.23-0.67 (mean 0.43) in France (Bacles et al., 2018). As in most cases, this reduction of *N*_*b*_ relative to census size primarily reflects skewed sex ratios and variance in reproductive success, even under controlled conditions. However, the absolute values of the Saimaa salmon effective population size remain particularly low across all scenarios, despite the species’ long generation time and the recent increase in the number of breeders. The main driver of this persistently low *N*_*e*_ appears to be the extremely small number of breeders during the early years (1980s–1990s) of the program, which likely caused a severe genetic bottleneck. A comparable pattern has been reported in a Sockeye salmon population maintained by a captive breeding program, where an effective population size of 41 individuals was estimated due to a critically low number of breeders at the onset of the program (Kalinowski et al., 2012).

Our study emphasizes the general importance of management practices and accurate records in captive breeding programs and has relevance for designing future programs. Estimates of the effective number of breeders (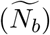) and effective population size *N*_*e*_ were sensitive to assumptions regarding variance in reproductive success. In particular, family-correlated survival reduced *N*_*e*_ by 55%, highlighting the strong influence of family-level demographic processes on the long-term genetic health of this population. Although this scenario may represent an extreme case - assuming that only half of the families survived - it illustrates the potential magnitude of such effects. In an empirical study on hatchery coho salmon, family-correlated survival has been shown to reduce *N*_*e*_ by approximately 20% (Moyer et al., 2007). Such effects could naturally occur in the Saimaa salmon system through family-dependent variation in post-release survival in the lake, a process that is difficult to mitigate through management measures. Nonetheless, our results also highlight the important role of controlled captive breeding practices in reducing reproductive variance at early life stages. The fertilization matrix system used at the Enonkoski hatchery, in which eggs from each female are divided and fertilized by several males (typically a 5×5 matrix design), and the equalization of family sizes at the eyed-egg stage, effectively minimize early-stage variance in reproductive success. Maintaining such practices is thus essential for maximizing *N*_*e*_, which increased by 12% in our scenario. In contrast, mass fertilization methods - where eggs and milt are mixed in bulk - can lead to highly unequal reproductive success among males and females and should be avoided. More generally, equalizing family sizes, even at the cost of discarding some offspring to match the smallest families, remains an effective measure to prevent disproportionate genetic contributions.

Other conservation measures are very important in addition to the captive breeding program. Despite its benefits, captive breeding alone cannot fully compensate for the limited number of breeders which constrains the effective population size of the Saimaa population. The increase in *N*_*e*_ over time reflects improvements in breeding stock management, the total ban on fishing of wild salmon (i.e., salmon with an adipose fin) implemented in 2016, and possibly higher survival rates in the lake. However, further increases in effective size will require continued efforts to increase the number of breeding individuals and allow natural reproduction. Since 2013, habitat restoration efforts and experimental releases of mature individuals into potential reproductive areas have yielded promising evidence of natural reproduction and juvenile recruitment (Leinonen et al., 2020; Hatanpää et al., 2021).

Nevertheless, even under optimistic scenarios, the effective size of this population remains below the common minimum 50/500 rule for short- and long-term persistence of closed populations (Franklin, 1980). This suggests that demographic recovery alone may not be sufficient to restore genetic diversity without additional interventions. Therefore, continued genetic monitoring of the population is essential. In addition, strategies such as assisted gene flow or genetic rescue could be considered. Experimental hybridization studies between Saimaa salmon and other Atlantic salmon populations (Eronen et al., 2021, 2024) have reported no evidence of outbreeding depression after one generation, and higher pre- and post-hatching survival in hybrids compared to pure-bred land-locked salmon (Saimaa). However, Klemme et al. (2024) found more contrasting results, and the effects of hybridization need to be evaluated over the entire life cycle and in backcross generations, where recombination could reveal hidden incompatibilities. More generally, in very small populations such as this one, caution is warranted in genetic rescue, as introduced deleterious variation may interact with inbreeding load and be difficult to purge, with potential delayed fitness effects (Pérez-Pereira et al., 2022).

While our demographic framework simplifies some aspects of the captive-breeding program, it provides valuable insights into the range of plausible long-term *N*_*e*_ values of this endangered population. Some assumptions may lead to overestimation of *N*_*e*_, such as equal reproductive success among males, constant variance in reproductive success across generations, whereas others may result in underestimation, including the assumption of strong family-correlated mortality. The model also assumes semelparity, although Atlantic salmon can be iteroparous and some individuals may reproduce multiple times across breeding stock cycles. In addition, the captive-breeding program is structured around continuous breeding stock lines composed of overlapping age classes (typically three to four). This design helps prevent close inbreeding by mating individuals from different cohorts but does not substantially increase long-term *N*_*e*_, as all breeding stocks ultimately trace back to a small number of wild founders.

In conclusion, our findings highlight several general principles for the management of captive breeding and release programs. First, maximizing the number of wild breeders and minimizing variance in reproductive success are key to maintain effective population size. Second, family-size equalization and monitoring of post-release survival can help mitigate family-correlated mortality. Demographic-based estimates, such as those presented here for the Saimaa salmon, can provide valuable tools for assessing the genetic risks of small, managed populations. Applied to the Saimaa salmon, our results indicate that, despite decades of captive breeding and release, the population remains extremely vulnerable. Continued genetic monitoring, careful breeding stock management, and restoration of natural reproduction are urgently needed to preserve the long-term evolutionary potential of this unique landlocked lineage.

## Acknowledgments

We thank the staff of Enonkoski Aquaculture Station of the Natural Resources Institute Finland for the information on the hatchery practices. Marko Luhtala is thanked for suggesting the original idea. We also thank Aslak Eronen for helping with designing the map of the study system. This study was funded by the European Union (ERC, FishLEGs, 101054307). Views and opinions expressed are however those of the author(s) only and do not necessarily reflect those of the European Union or the European Research Council Executive Agency. Neither the European Union nor the granting authority can be held responsible for them. The authors declare no conflict of interest.

## 6 Data availability

Data and code used are available at https://zenodo.org/records/18496177?token=eyJhbGciOiJIUzUxMiJ9.eyJpZCI6IjcyYmEwNDg2LWE2NjQtNDRkMS04MDEyLTc1OTQ1Y2Y0OTNlYiIsImRhdGEiOnt9LCJyYW5kb20iOiIyMWYxNW mb37D6T7ASDoSrYKkT_wISHIEPd6ktzDh7aWTJKbl1tu6u2ewBs0WIa80f84YzYxNEYyh0HTH5qSEay7S0Cpzg.

## Notes

### Competing Interest Statement

The authors have declared no competing interest.

