## Supplementary Table S1 for "Extremely low effective population size in a captive-bred population: partial mitigation through management practices"

### Appendix S: Supplementary Materials

Table S1: Age distribution and generation length of wild-caught Saimaa salmon breeders by sex. Age at offspring birth corresponds to the time elapsed between hatching of a parent and hatching of its offspring. Generation length ($g$) was calculated as the weighted mean age at reproduction for females, males, and overall.

| | | **Female** | | **Male** | |
|---|---|---|---|---|---|
| Breeding age | **Age at offspring birth** | $N$ | Proportion | $N$ | Proportion |
| 4+ | 5 | 53 | 0.264 | 2 | 0.025 |
| 5+ | 6 | 85 | 0.431 | 30 | 0.37 |
| 6+ | 7 | 48 | 0.244 | 35 | 0.432 |
| 7+ | 8 | 12 | 0.061 | 14 | 0.173 |
| **Generation length ($g$)** | | **6.10** | | **6.75** | |
| | | **6.43** | | | |
